# Repetitive and Brief Occlusion of the Majority of the Vena Cava Flow Attenuates Pulmonary Arterial Hypertension and Improves *I*_K+_ of the Pulmonary Arterial Smooth Muscle Cells in Rat Models: a novel potential therapeutic strategy for PAH

**DOI:** 10.1101/2020.10.04.314708

**Authors:** Nan Liu, Ying Xing, Lv Wang, Chen Wang

**Affiliations:** department of Geriatrics, Xijing Hospital, the Fourth Military Medical University, Xi’an, China; department of Cardiology, Xijing Hospital, the Fourth Military Medical University, Xi’an, China; Clinic of Cardiovascular & Metabolic Disease, W.X.Y.Z.-Xi’an medical center, Xi’an, China; department of Endocrinology, Xi’an Daxing Hospital, Xi’an, China; department of Thoracic Surgery, Xi’an Daxing Hospital, Xi’an, China

**Keywords:** Treatment, Hemodynamics, Potassium Current

## Abstract

By now, there are limited therapeutic approaches satisfactory for pulmonary arterial hypertension (PAH) treatment. In the present study, we established a novel rat model, repetitive and brief occlusion (RBO) of the majority of the right atrium venous blood flow in the conscious rats. Using this model, we found that the RBO treatment significantly improved the pulmonary hemodynamics with a lower pulmonary vascular resistance observed. The I_K+_ and big-conductance Ca^2+^-activated K^+^ (BK) channels were involved in the mechanism of the RBO treatment for PAH. Although the results are limited, the conclusion are solid and we believe this manuscript is valuable for all the researchers and clinical doctors who are interested in PAH.

---

Pulmonary arterial hypertension (PAH) is a cardiovascular disorder associated with the severe morbidity and mortality. The pathophysiology of PAH remains controversial, and the current available therapies for PAH are limited and not satisfactory. Chronic pulmonary artery (PA) overflow could induce PA injury and PAH in animal models and in patients, and hemodynamic unloading by PA banding could prevent or even reverse distal PA lesions in PAH models [1], which indicated hemodynamic stress was an important factor in the development of the PAH with vascular structural injury and the high vascular resistance. It is suggested that a feedback circle of PAH exists: overflow or mechanical stretching of the PA vascular wall triggered the PA constriction or occlusion, inducing the high vascular resistance and the high pressure in PA, and then the higher pressure in PA further again fed back to vascular wall, increasing more mechanical stress and stretching, and again. The real PAH should include the whole positive feedback circle. Therefore, we asked whether a stimulus of low flow and low pressure could break the feedback circle and could reverse the high vascular resistance or PAH with dilation of the PA?

To elicit the stimuli of low flow and low pressure on pulmonary arterial wall, we established a novel model: repetitive and brief occlusion (RBO) of both anterior vena cava (AVC) and posterior vena cava (PVC) in conscious rats. In brief, two day before RBO treatment, adult male SD rats were under anesthesia with pentobarbital sodium (30 mg/kg, IP) and inhalation anesthesia with isoflurane. One transverse incision was made along the clavicles, then mobilization of both sides internal and external jugular veins by blunt dissection. And then the forelimbs were restricted together and pulled to cranial side, which posture was to prevent the injury of brachial plexus in next procedure. A 2-0 suture with 25mm round needle were inserted and wrapped the internal and external jugular veins and then penetrated out of the subclavian space (same for both sides), which operation could totally wrap the anterior vena cava (internal jugular vein, the external jugular vein and the subclavian vein, Fig1A). The ends of all threads were outside the body of the rat, and then muscle and skin layers were closed. Next, make a transverse dorsal incision along the 9th or 10th intercostal space and develop with a blunt hemostatic forceps until retroperitoneal fat was seen. A cotton swab was used to develop the retroperitoneal space for the forceps insertion between the liver and the diaphragm. Subsequently, mobilization of PVC by blunt dissection and a hook was introduced gently into the retroperitoneum and took the PVC close to the intercostal space and then inserted a 1-0 suture and suspended the PVC then released and returned the PVC to the retroperitoneal space. Closed the muscle and skin layers and the ends of the suture were left outside of the rat body. Therefore, for both AVC and PVC, we could make the threads into an overhand knots outside the body and the ligatures could be tightened and loosened to get the RBO treatment in conscious rats. The same surgical procedures were carried out in the sham groups, but loose knots were made and loosened during sham treatment. After RBO treatment, all threads were cut and pulled out slowly. During the RBO treatment, we repetitively and briefly occluded both AVC and PVC by ligation (occlusion for less than 5 seconds then re-open for 30 seconds and repeated 5 cycles as one sequence, 1 sequence every 6 hours) to intermittently restrict right ventricular (RV) preload, for continuous 24 hours, total 5 sequences, in the Sugen 5416 (VEGF receptor blocker) and hypoxia induced PAH rat model [1].

**Fig. 1.**
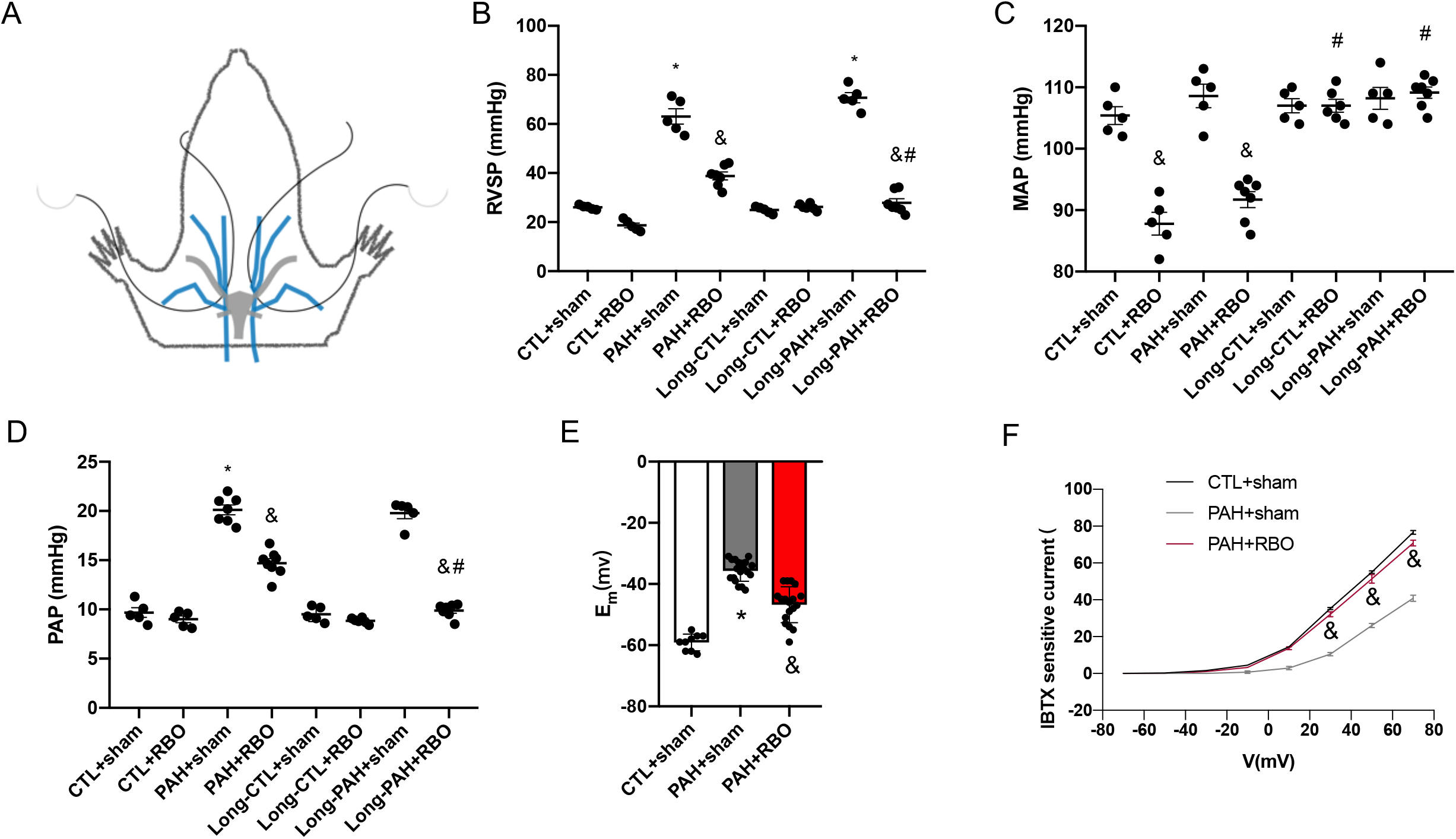
RBO treatment improves RVSP in vivo and PAP in vitro. A, the schematic diagram for the prepare of AVC; B,C, in vivo study; D, isolated lungs perfusion study; E,F, cell electrophysiological study. * p<0.01 compare to CTL, & p<0.01 compare to sham, # p<0.01 compare to acute group. RVSP: right ventricular systolic pressure; MAP: mean artery pressure; PAP: pulmonary artery pressure.

To assess the acute effects of RBO treatment, hemodynamic evaluation was performed after 1 hour rest following the 24-h-RBO treatment. In studies of acute effects of RBO in vivo, the mean peak RV systolic pressure (RVSP) was significantly lower in PAH+RBO group (39.1±1.1 mmHg, n=7) than in PAH+sham group (62.2±2.9 mmHg, n=5), shown in Fig.1B. In the PAH+RBO group, mean artery pressure (MAP) decreased significantly 14% (Fig.1C) and cardiac output (CO) decreased significantly 4% compared to the PAH+sham group. To directly examine the acute effect of RBO on the pulmonary vascular resistance (PVR) in lungs, we isolated lungs and performed perfusion experiments (Cadogan, 1999) after RBO treatment finished in vivo. Lungs were perfused with a mixture of blood and physiological saline solution. Perfusion was maintained at a constant flow (0.04 ml·min^-1^·Kg^-1^) so that changes in arterial perfusion pressure reflected changes in total PVR. Mean pulmonary arterial pressure (PAP) in PAH+sham group (n=7) was 20.3±0.7 mmHg and was significantly reduced to 14.2±0.8 mmHg in PAH+RBO group (n=8, Fig. 1D).

To observe the long-term effects of the RBO treatment, we measured hemodynamics in vivo after 1 week refeeding following the RBO treatment in independent groups. The mean peak RVSP in long-term PAH+RBO group (Long-PAH+RBO) deceased further (27.3±1.7 mmHg, n=7, Fig. 1B) than in the acute PAH+RBO. The acute reduction of MAP and CO in PAH+RBO group observed above was largely reversed in Long-CTL+RBO and in Long-PAH+RBO group (Fig. 1C). We also measured the PAP in isolated lung perfusion experiment in the long-term groups. PAP in Long-PAH+RBO group was 9.7±0.8 mmHg (n=7, Fig. 1D), which was significantly lower than in Long-PAH+sham group and lower than the acute effect in the PAH+RBO group. Therefore, we confirmed additional beneficial effects of RBO treatment after a period of normal feeding, including further decreasing in PVR.

To find out the possible mechanisms of the RBO treatment, we did the cell electrophysiological studies. We know the potassium currents (*I*_K+_) in the pulmonary vascular smooth muscle cells (PASMCs) play an important role in the pulmonary artery dilation and constriction [2]. Inhibition of *I*_K+_ leads to membrane depolarization and Ca^2+^ entry through the voltage-dependent Ca^2+^ channels, which is one of mechanisms of vascular constriction and PAH (Weir, 1995;1996). Big-conductance Ca^2+^-activated K^+^ (BK) channels are highly expressed in PA smooth muscle cells and play a critical role in regulating the resting membrane potential. Moreover, BK currents were inhibited in PAH models, and activation of BK channels could hyperpolarize the membrane of the vascular smooth muscle cells and dilate the vessels (Nelson, 1995), which was proven effective in attenuating PAH. Furthermore, BK was reported mechanosensitive (Kirber, 1992; Zhang, 2018). So, we hypothesized that the inhibition of BK currents in PASMCs in PAH rat model was partially reversed by the RBO treatment. To test our hypothesis, we did the electrophysiological studies on freshly dissociated rat PASMCs as previously described [3]. The resting *E*_m_ of PASMCs in PAH+RBO group (n=19) was more polarized than in PAH+sham group (n=18) (Fig.1E) as expected. The whole-cell *I*_K+_ of BK channels (IBTX sensitive currents) in PASMCs was largely decreased in isolated PASMCs of PAH+sham group and totally reversed by RBO treatment as shown in Fig.1F.

The restriction or reduction of RV preload might be previously thought as an adverse effects by medicine treatment for PAH because of the possible systemic hemodynamic deterioration. In the present study, however, we found this RBO treatment was beneficial for lowering PVR and improving PAH, which provided a potential therapeutic strategy for PAH in human. We would like to name this as a hypo-load induced vasodilation phenomenon.

While the mechanical stimuli on vascular smooth muscle were studied widely, the most of them focused on the positive pressure and stretching mechanical stimuli but the low pressure mechanical stimuli on vascular wall were not clear. However, the thoracic cavity and lungs were the negative pressure regions. So, understanding of the low pressure mechanical stimuli on the PASMCs is important. The characters of the present RBO rat model included: briefly stopped majority of RV venous supply; left heart keeping pumping and draining the pulmonary vascular content; the spontaneous breathing (the negative pressure driven breathing) going on; intact vascular structure in vivo but not isolated cells or vascular rings in vitro. All of these factors composed the low pressure mechanical stimuli on PA.

The mechanism of effects of RBO might be that the low pressure induced PA vascular wall loose and collapse which, as mechanical stimuli, improved *I*_K+_ by activation of the BK channels and hyperpolarized the membrane of PASMCs and reduced the Ca^2+^ entry and then decreased the PVR and attenuated the PAH. Membrane depolarization was also involved in proliferation of PASMCs (Platoshyn, 2000), and RBO treatment could inhibit the depolarization of PASMCs in rat models. So it was possible that the reversed pulmonary vascular medial hypertrophy was involved in the mechanisms of the additional benefits in Long-term RBO group. These findings were consistent with the results that hemodynamic unloading by pulmonary artery banding nearly fully reversed the established occlusive pulmonary arterial lesions and perivascular inflammation in advanced PAH rat models [1].

In some studies, intermittent occlusion of AVC or PVC alone by the balloon in swine models also elicited the hemodynamic change and got some cardioprotective results (Kapur, 2020), but in our earlier preliminary tries, intermittent occlusion of AVC or PVC alone was not enough to get positive effects after the 24-h treatment in rats. But we don’t exclude the possibility that positive results might also be appear if we prolonged the treatment time and added the sequence times in rat models. Limits of the present study: this is an early and preliminary study and we only provided the functional results. We did some molecular biological and histochemical studies but didn’t get positive results by now, so we are trying to do more models and to observe a longer-time result.

